# KG-COVID-19: a framework to produce customized knowledge graphs for COVID-19 response

**DOI:** 10.1101/2020.08.17.254839

**Authors:** Justin Reese, Deepak Unni, Tiffany J. Callahan, Luca Cappelletti, Vida Ravanmehr, Seth Carbon, Tommaso Fontana, Hannah Blau, Nicolas Matentzoglu, Nomi L. Harris, Monica C. Munoz-Torres, Peter N. Robinson, Marcin P. Joachimiak, Christopher J. Mungall

## Abstract

Integrated, up-to-date data about SARS-CoV-2 and coronavirus disease 2019 (COVID-19) is crucial for the ongoing response to the COVID-19 pandemic by the biomedical research community. While rich biological knowledge exists for SARS-CoV-2 and related viruses (SARS-CoV, MERS-CoV), integrating this knowledge is difficult and time consuming, since much of it is in siloed databases or in textual format. Furthermore, the data required by the research community varies drastically for different tasks - the optimal data for a machine learning task, for example, is much different from the data used to populate a browsable user interface for clinicians. To address these challenges, we created KG-COVID-19, a flexible framework that ingests and integrates biomedical data to produce knowledge graphs (KGs) for COVID-19 response. This KG framework can also be applied to other problems in which siloed biomedical data must be quickly integrated for different research applications, including future pandemics.

**BIGGER PICTURE:** An effective response to the COVID-19 pandemic relies on integration of many different types of data available about SARS-CoV-2 and related viruses. KG-COVID-19 is a framework for producing knowledge graphs that can be customized for downstream applications including machine learning tasks, hypothesis-based querying, and browsable user interface to enable researchers to explore COVID-19 data and discover relationships.

## INTRODUCTION

Although most coronaviruses typically cause common-cold symptoms in humans, three betacoronaviruses have emerged in the last few decades that can cause a range of serious manifestations including pneumonia and death: the severe acute respiratory syndrome (SARS) coronavirus (SARS-CoV-1), the Middle East respiratory syndrome coronavirus (MERS-CoV), and the novel betacoronavirus that emerged in late 2019, subsequently named SARS-CoV-2, the agent of the disease COVID-19.^1^ The rapid spread of SARS-CoV-2 has led to a global pandemic.

COVID-19 is a complex disease involving many biological processes and pathways, each of which involves many genes. Initial symptoms of COVID-19 typically include fever, cough, fatigue, anorexia, anosmia, myalgia, and diarrhea. In some patients, severe illness ensues roughly one week after the initial onset of symptoms, and can present with rapidly progressive respiratory failure.^2^ In addition to the symptoms highlighted, COVID-19 infections can lead to secondary health problems such as blood clots^3^, tissue necrosis, organ damage, and, in some cases, cardiac failure. Given that the research community is still learning about COVID-19, its symptoms and their underlying pathological mechanisms are still being uncovered.

Many possible treatments for different aspects and stages of COVID-19 are being actively pursued. Evidence suggests that remdesivir (DrugBank:DB14761) can shorten the time to recovery in adults hospitalized with COVID-19 infection and pneumonia (though the effect is not statistically significant),^4^ and more recent evidence suggests that dexamethasone (DrugBank:DB01234) may reduce mortality in patients with severe COVID-19.^5^ However, currently no treatment is available to prevent progression of COVID-19 to severe disease, and our knowledge of the causes and optimal medical management of the many clinical complications of COVID-19 is limited.

A large amount of biomedical and molecular data is available to aid the massive research effort to address the COVID-19 pandemic. Before the pandemic began, there existed a large amount of biomedical data for coronaviruses other than SARS-CoV-2 (SARS-CoV and MERS-CoV^6^ as well as many other pathogenic and non-pathogenic coronaviruses), such as viral genome and transcriptome sequences, viral/host gene interactions, gene function, epidemiological data, and clinical case data. Much of this information is now also available for SARS-CoV-2. In addition, there is also a large amount of data about drugs that may offer a treatment for COVID-19, as well as the protein targets for each drug.

However, researchers are confronted with a number of technical challenges when trying to use existing data to discover actionable knowledge about COVID-19. The data needed to address a given question are typically siloed in different databases and employ different identifiers, data formats, and licenses. These data sources are often in different formats, requiring transformation in order to serve the task at hand. For example, to examine the function of proteins targeted by FDA-approved antiviral drugs, one must download and integrate drug, drug target, and FDA approval status data (from Drug Central, for example, in a bespoke TSV format^7^) and functional annotations (from, for example, Gene Ontology in GPAD format^8^). Furthermore, many data sets are updated periodically, which requires researchers to re-download and re-harmonize data in order to perform their analysis on the most current data.

To tackle the daunting challenge of bringing together these disparate sources of information and extracting useful knowledge from them, we employed knowledge graphs (KGs). Knowledge graphs are a way to represent and integrate heterogeneous data and their interrelationships. In a KG, discrete entities or pieces of information form distinct nodes interconnected by edges, where both nodes and edges are typed using a hierarchical system such as an ontology^9^.

For example, nodes of type *‘protein’* representing individual entities (such as human ACE2 or SARS-CoV-2 Spike) can be interconnected via edges of type ‘*orthologous to*’ or ‘*interacts with*’, and these nodes can be connected with other kinds of nodes representing diseases, drugs, and so on. This kind of representation is amenable to complex queries (e.g. “which drugs target a host protein that interacts with a viral protein?”), and also to graph-based machine learning (ML) techniques.

## RESULTS

### The KG-COVID-19 Framework

We created KG-COVID-19 to address the challenge of integrating data for COVID-19 response. KG-COVID-19 is a framework that enables the creation of customized KGs containing COVID-19 knowledge for different applications. For example, a drug repurposing application would make use of protein data linked with approved drugs, while a biomarker application could utilize data on gene expression linked with pathways. The methodology is not limited to COVID-19, but could support data integration for any biomedical research effort.

### Constructing the knowledge graph

Our process for generating the KG was designed to support interoperability, preserve provenance, and provide the ability to flexibly mix and match data from different sources. The workflow is divided into three steps: data download (fetch the input data), transform (convert the input data to KGX interchange format), and merge (combine all transformed sources) (Figure 1).

**Figure 1.**
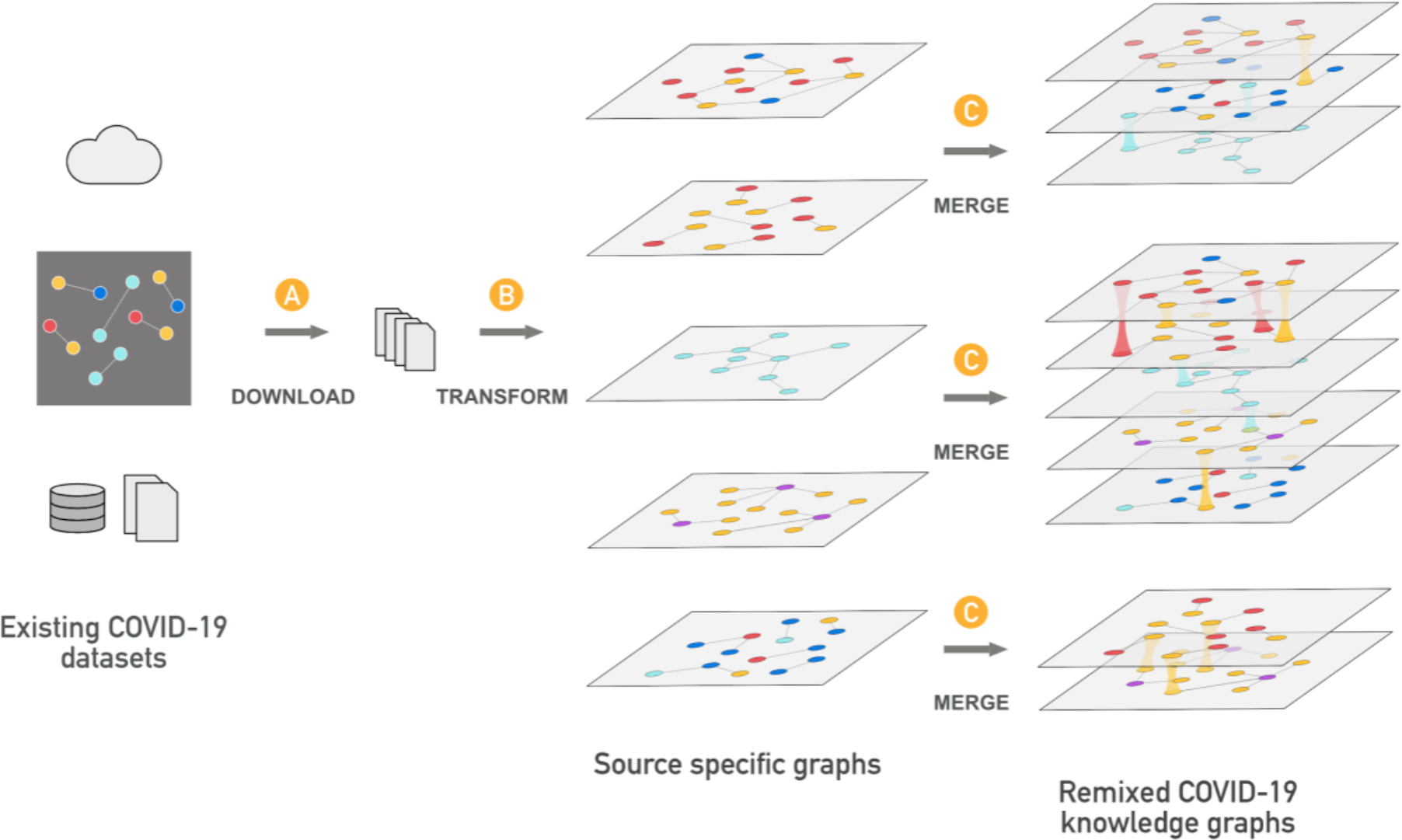
The KG-COVID-19 framework for producing KGs. The framework is divided into three modular steps: download, transform, and merge. A) The download step retrieves all data sets needed for ingestion using a set of URLs specified in a YAML file. B) The transform step applies Python code that is specific to each source to transform the most useful elements of each source and emit a graph in TSV format. C) The merge step uses a YAML file to read the user-specified data sets (among those produced in the transform step) and merge them into a single KG. Different YAML files can be constructed to mix and match different input data from B, but each merge operation yields a single merged graph. Both the transform and merge steps rely heavily on KGX, a powerful tool for manipulating knowledge graphs (https://github.com/NCATS-Tangerine/kgx).

#### Download

The download step retrieves data from multiple sources using a YAML file that specifies the source URLs (Figure 1A). Our experience has shown that this step is a frequent point of failure in many extract, transform, and load (ETL) pipelines and separating out this step helps mitigate this issue.

The data sources we ingest are focused on our use case: drug repurposing (e.g., drug and drug target data, protein interaction data, ontologies important in disease such as HPO and Mondo). However, we also ingest data sources that our user community requests by opening tickets on our project GitHub page.^10^

#### Transform

The transform step (Figure 1B) involves parsing the input files and translating them to a graph-based representation. We have devised a simple yet expressive format called KGX interchange format^11^ - a serialization for representing a graph that combines features of resource description framework (RDF) and property graphs. KGX interchange format consists of two tabular files, one for representing graph nodes and their properties, the other for representing edges and their properties (Figure 2). Using standards from the semantic web, nodes in the graph are identified by Compact Uniform Resource Identifiers (CURIEs).^12^ These can be expanded to an Information Resource Identifier (IRI), which is the global identifier for this node. All nodes are assigned a type using the *‘category’* node property, and all edges are typed using the *‘edge_label’* property. Where possible, one can use classes from the Biolink Model,^13^ a high-level data model for representing biological and biomedical knowledge. Granular typing of nodes is possible by adding additional classes to the *‘category’* property. Granular typing of edges is possible by adding a more specific relation to the ‘*relation’* property. For example, one can use a class from the Relation Ontology (RO)^14^ to further classify the semantics of an edge.

**Figure 2.**
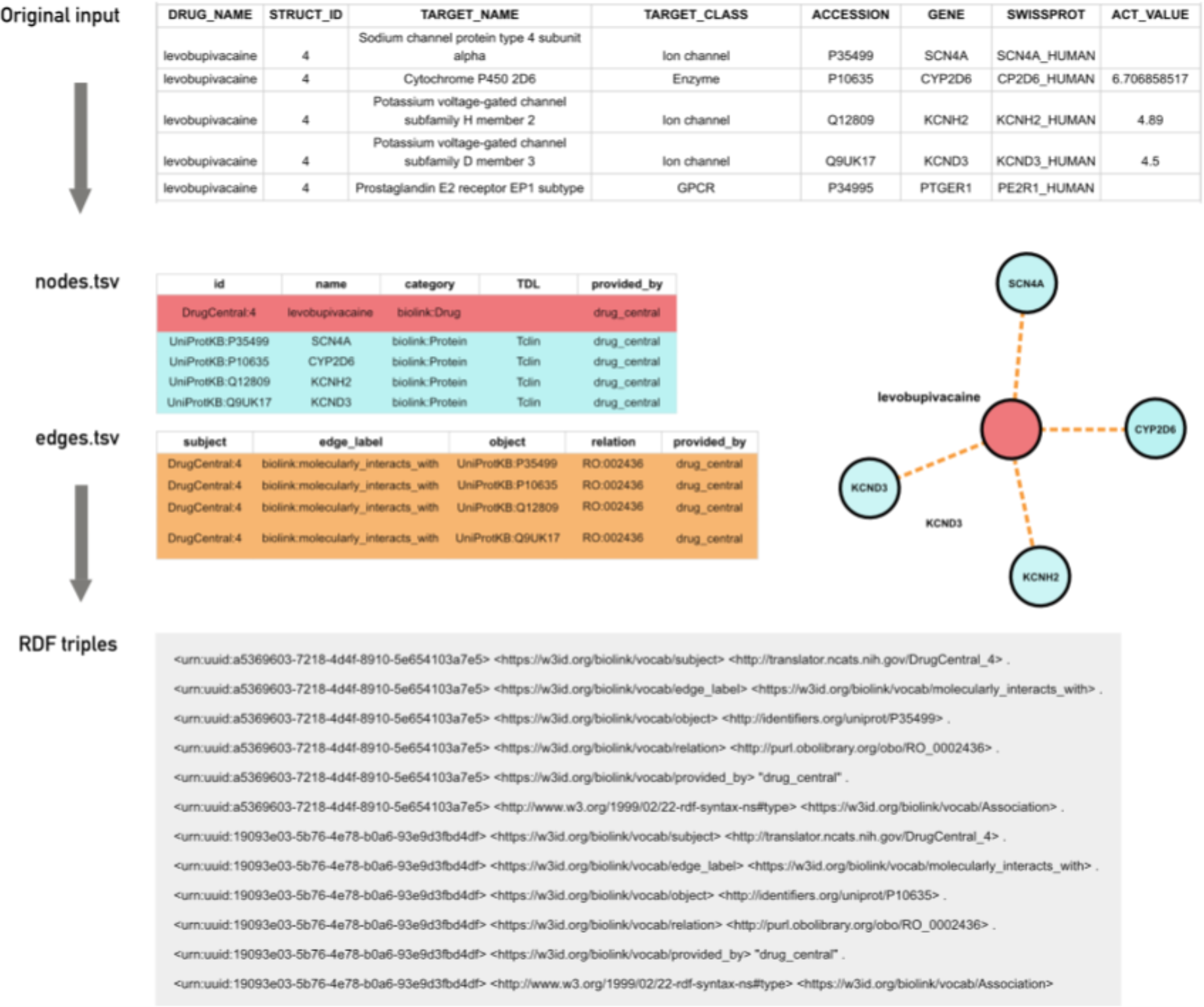
A typical transformation of records from an input file into entries in a nodes.tsv and edges.tsv file representing the nodes and edge in a graph. These nodes and the edge can be further transformed into RDF triples.

#### Merge

The merge step (Figure 1C) combines the component data sets into a KG. This step is informed by a YAML file that specifies what data sets should be included, to allow for flexible remixing of subgraphs. In addition to selecting different component data sets to be merged, the user can also filter nodes and edges from each source by the node ‘*category’* and ‘*edge_label’*, allowing fine grained control of the resulting graph. By default, all nodes and edges from all component data sets are merged. Optionally, the merged graph can be loaded into any triple/RDF store or Neo4j database.

### Design principles

While our framework offers flexibility in deciding how best to transform each data source, KG-COVID-19 follows some general design principles to maintain the quality of the resulting KG.

#### Ensure interoperability through standardized node and edge representations

We use a core set of standardized ontologies and the Biolink Model,^13^ a biological data model for categorizing nodes and edges, to facilitate interoperability and data summarization. To ensure Biolink Model compliance, a Biolink category and a Biolink predicate are required for the categorization of nodes and edges, respectively. Since Biolink predicates are typically very broad in scope, the edge can be further categorized by adding a more specific description in the *‘relation’* property using a term from the Relation Ontology.^15^ Categorization using ontologies and the Biolink Model provides a convenient way to assess what types of data have been ingested from each source, record provenance information, and also facilitates interoperability with other transformed data sets.

#### Ingest only relevant data

Only the subset of features in each data set that are likely to be useful downstream are preserved, and only statements for which the source is authoritative are ingested (for example, assertions about protein interactions are not ingested from a drug database).

#### Normalize identifiers at the time of ingest

Identifier (ID) normalization is crucial for ensuring connectedness and the utility of the graph. We refer to the Biolink Model to provide the preferential order of identifier prefixes to be used for a particular Biolink class. For example, in the case of Gene class (https://biolink.github.io/biolink-model/docs/Gene) the model prescribes HGNC, NCBIGene, ENSEMBL, where the order of prefixes matters: identifiers from HGNC namespace are given a higher priority than NCBIGene and ENSEMBL. In the case of Protein class, the model prescribes UniProtKB identifiers. For drugs and other chemical compounds, the model recommends the following: CHEBI, CHEMBL, DrugBank, PubChem. Identifiers can also be normalized by adding cross-references to other identifiers in the *‘xrefs’* property of nodes, which is the *‘xrefs’* column in the KGX interchange format TSV describing the nodes.

#### Preserve provenance

Each ingest adds a ‘*provided_by’* column in the edge TSV file, which ensures that graphs into which the data are merged (Figure 1C) contain a record of which ingest produced each edge. The preservation of all files used to generate the graph in the download step (Figure 1A) makes it possible to trace each node and edge to the entries in the input file that generated them. PubMed IDs are added to the *‘publication’* column, where available, to provide additional provenance.

### Downstream tooling for querying and machine learning

The KG-COVID-19 framework contains tooling for common graph operations. The framework can create training and test data sets in graph form for machine learning applications such as training classifiers or regressors for link prediction (see Experimental Procedures). It also includes a query function that can execute prewritten or custom SPARQL queries on a given SPARQL endpoint (by default, our endpoint: http://kg-hub-rdf.berkeleybop.io/blazegraph/#query).

### Current contents of KG-COVID-19

A schematic diagram of all data sources currently ingested is shown in Figure 3. The data we ingest are focused on sources relevant to drug repurposing for our downstream querying and machine learning applications, prioritizing drug databases, protein interaction databases, protein function annotations, COVID-19 literature, and related ontologies. The KG contains drug and chemical compound data from several databases (currently DrugCentral,^16^ the Pharmacogenomics Knowledgebase (PharmGKB),^17^ Therapeutic Target Database (TTD),^18^ and ChEMBL^19^), functional annotations and synonyms for coronavirus genes and proteins from the Gene Ontology (GO), and protein interaction data from STRING^20^ and the IntAct Molecular Interaction Database^21^. We ingest data about the occurrence in COVID-19 scientific publications of concepts such as Gene Ontology (GO) terms, UniProt Knowledgebase (UniProtKB) proteins, National Center for Biotechnology Information (NCBI) and HUGO Gene Nomenclature Committee (HGNC) genes, and ChEMBL IDs via SciBite annotations^22^ of the COVID-19 Open Research Dataset (CORD-19).^23^ To capture ontology-based annotations, the relational graphs for the GO,^8^ Human Phenotype Ontology (HPO),^24^ and Mondo Disease Ontology^25^ are ingested, and annotations are added to the graph as provided by each ingest.^26^ In addition, we ingest GO-CAM models that capture biological systems such as protein pathways, including those important in SARS-CoV-2 infection.^27^

**Figure 3.**
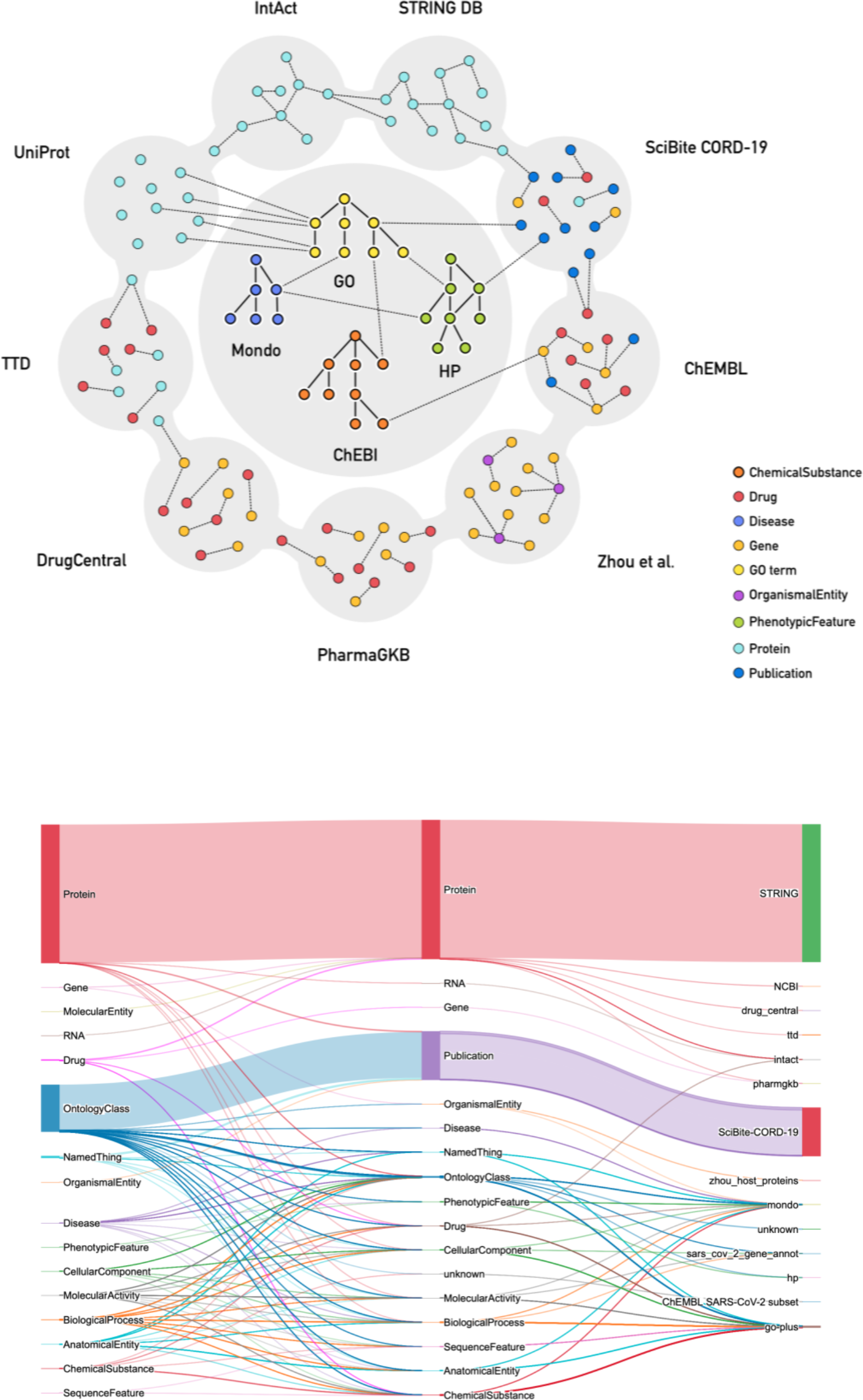
Schematic representation of the data currently ingested into the KG-COVID-19 knowledge graph. (Top) Polygons shown correspond to the various data sources currently ingested into the KG, and the small colored circles indicate the data types ingested from this source. (Bottom) Sankey plot showing the Biolink categories for edges in the KG-COVID-19 graph. Left and middle columns show Biolink categories for edges, right column indicates the source of the data from which the edges were derived. Line widths are proportional to the number of edges.

### Use cases

While we designed KG-COVID-19 to allow flexible reuse and remixing of data to produce custom KGs, our immediate use case is to provide a COVID-19 KG that can be used for machine learning to produce actionable knowledge about COVID-19 (Figure 4). This use case demonstrates several features of KG-COVID-19, namely: normalization and merging of disparate data sources with different namespaces and formats, flexible remixing of component subgraphs, and a regular update cycle to keep up with new knowledge. We follow the workflow described in Figure 1 to produce the KG-COVID-19 knowledge graph. From the final merged graph, KG-COVID-19 produces training and test data sets suitable for machine learning applications (see Experimental Procedures). Embiggen^28^ (paper in preparation), our implementation of node2vec and related machine learning algorithms, is applied to this KG to generate embeddings, vectors in a low dimensional space which capture the relationships in the KG. Embiggen is trained iteratively to identify optimal node2vec hyperparameters (walk length, number of walks, *p, q* etc.) and to then train classifiers (e.g., logistic regression, random forest, support vector machines) that can be used for link prediction. The trained classifiers can then be applied to produce actionable knowledge: drug to disease links, drug to gene links, and drug to protein links. The latter would indicate a drug that might be useful for COVID-19 treatment.

**Figure 4.**
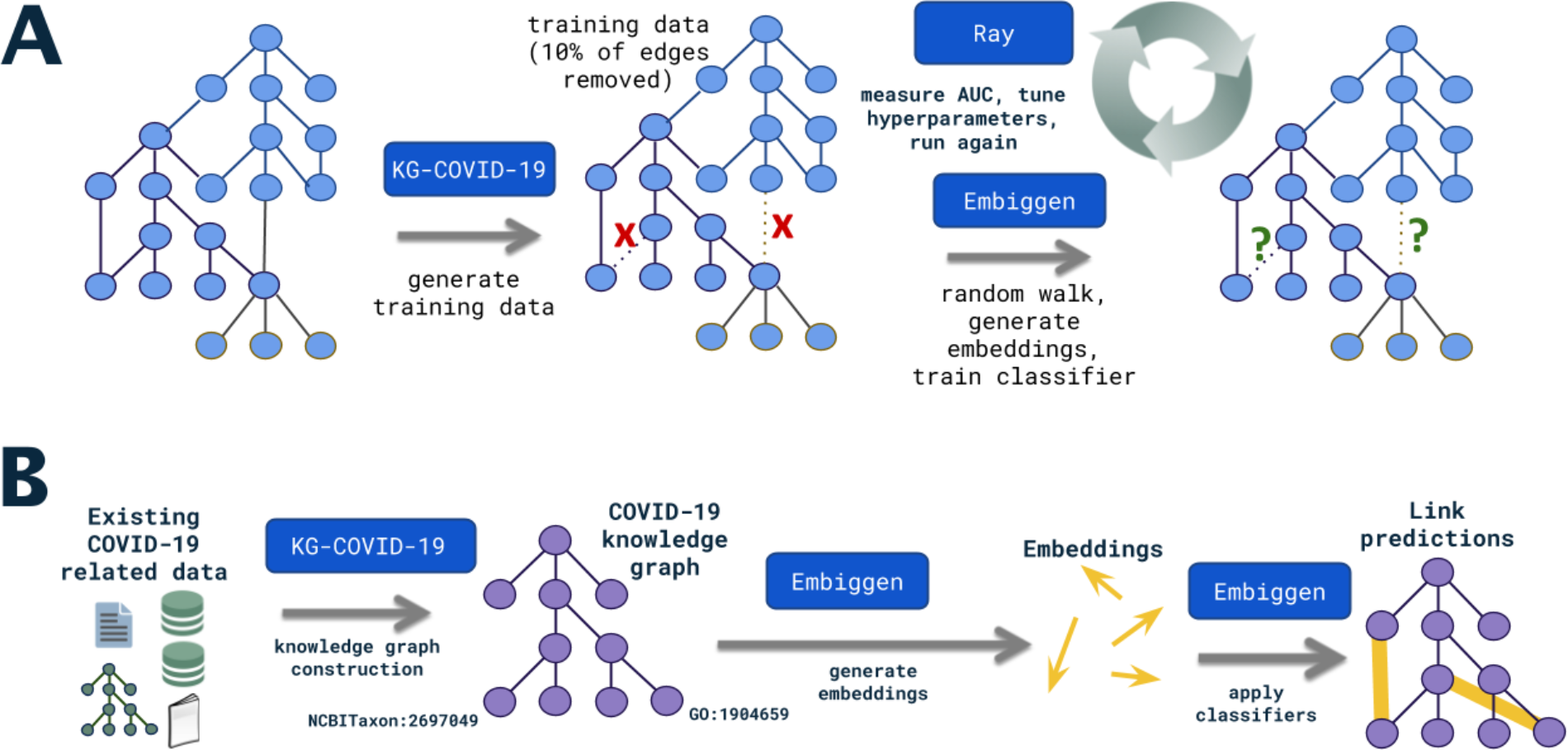
Workflow for machine learning application of KG-COVID-19 knowledge graph. A. In order to train classifiers for use in link prediction, training and test graphs are first produced from the original KG-COVID-19 graph (see Experimental Procedures). These graphs are used by Embiggen to generate random walks, embeddings, and finally a classifier. The test graphs are used to assess the performance of the classifier. This step is performed iteratively in order to identify optimal hyperparameters. B. The classifiers are applied to the KG-COVID-19 to perform link prediction in order to identify links that correspond to actionable knowledge: for example, links between drugs and the COVID-19 disease, links between drugs and SARS-CoV-2 protein targets, and links between drugs and host proteins that are involved in COVID-19 disease processes.

To demonstrate the usefulness of KG-COVID-19 for machine learning applications, we created embeddings for nodes and edges from the KG-COVID-19 knowledge graph and visualized the embeddings in two dimensions using a t-SNE plot (Figure 6). While only the graph structure and no biological typing of nodes was used to generate the embeddings, the nodes exhibited a tendency to cluster according to biological types. This indicates that the embeddings encode biological information that can be used for machine learning. Similarly, a t-SNE plot of edges in KG-COVID-19 displays grouping according to the type of the edge (Supplementary Figure 2).

**Figure 5.**
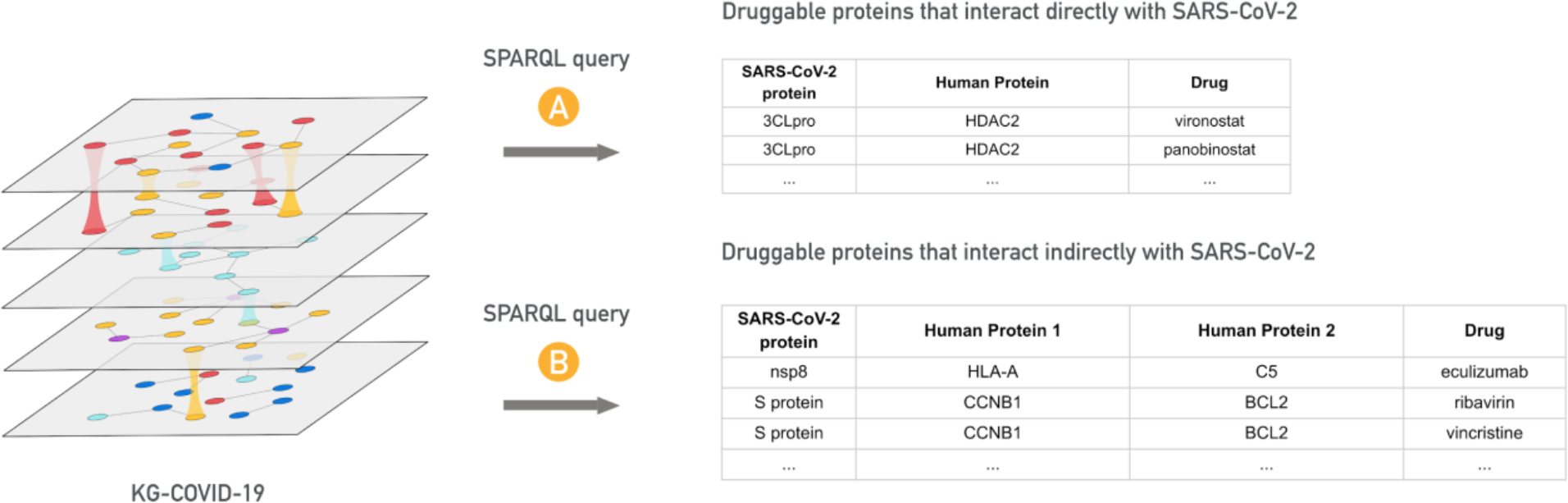
Hypothesis-based querying of KG-COVID-19 knowledge graph for using SPARQL queries. (Top) A SPARQL query retrieves approved drugs that target human proteins that physically interact with SARS-CoV-2 protein. (Bottom) A SPARQL query retrieves approved drugs that target human proteins that physically interact indirectly with SARS-CoV-2 through another human protein. The suitability of these drugs for repositioning are evaluated by NVBL collaborators, for example by analyzing available structural data to support repositioning.

**Figure 6.**
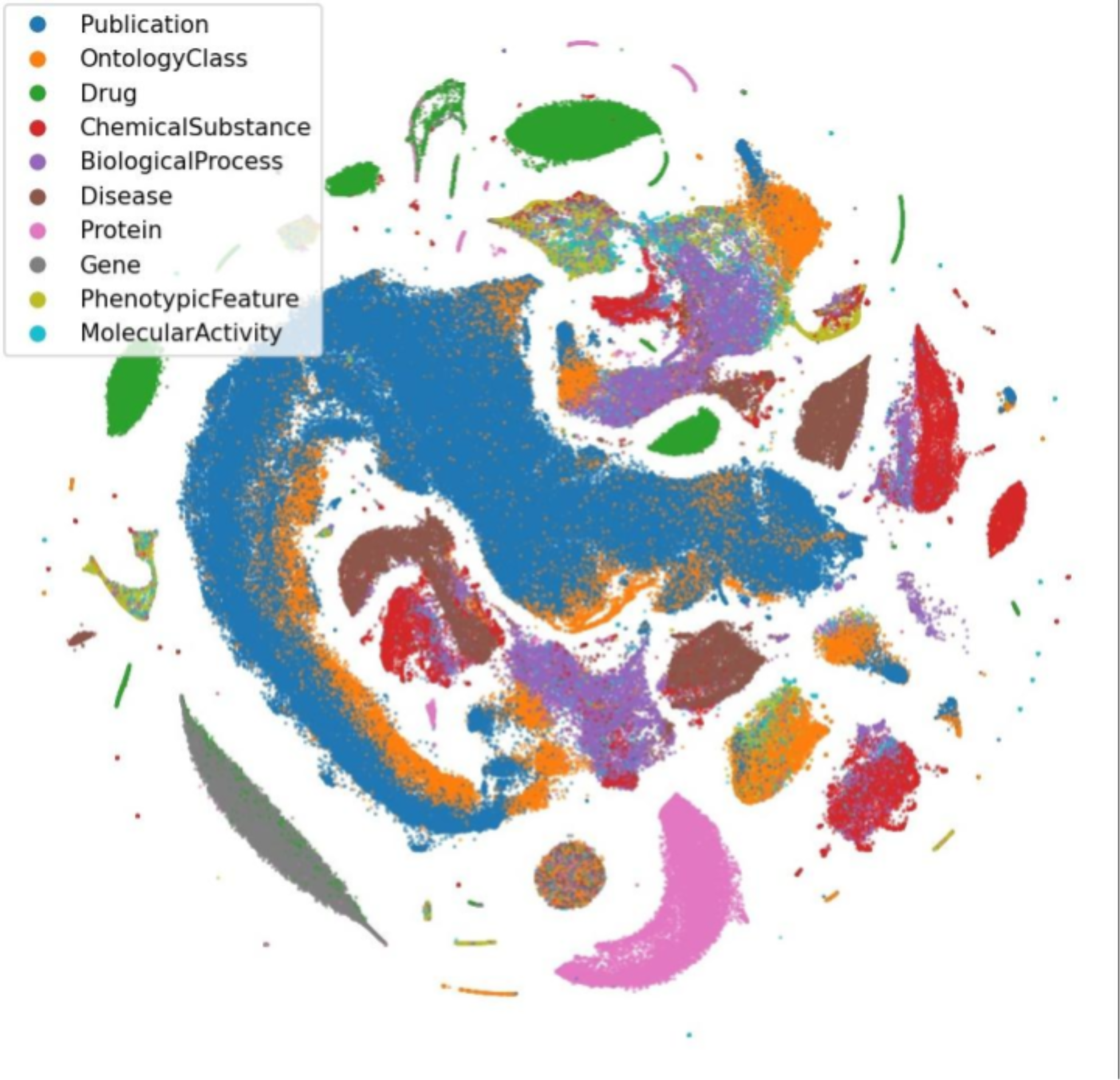
Visualization of KG-COVID-19 knowledge graph node embeddings using t-SNE. Embeddings were created for each node in the KG-COVID-19 knowledge graph and t-SNE was performed as described in Experimental Procedures. Nodes categorized with one of the ten most numerous Biolink categories were then selected. Colors indicate the Biolink category for each node.

While the initial development of KG-COVID-19 has focused on our machine learning applications, other use cases have emerged. As part of the National Virtual Biotechnology Laboratory (NVBL), we have found it useful to perform hypothesis-based querying of the KG to identify viral and human proteins that make attractive drug targets^29^. For example, we have queried the KG to identify host proteins that are known to interact with viral proteins, and these are further filtered according to whether these host proteins are targets of approved drugs, (Figure 5). These data are further analyzed with downstream analyses to assess their suitability for drug repurposing. Our KG is also part of a federated query used by the NVBL to collate and share up to date information related to COVID-19 and SARS-CoV-2. In addition, the National COVID Cohort Collaborative (N3C) has incorporated our KG as an ontologically-informed way to combine their clinical data sets (by virtue of our integration with GO, HPO and Mondo). The N3C also uses our KG to incorporate all of our transformed and harmonized data, saving them the onerous task of collecting and integrating all of those data sources individually.

## EXPERIMENTAL PROCEDURES

### KG generation pipeline

The framework to produce our KG is essentially an extract, transform, and load (ETL) system with additional tooling to facilitate downstream uses (e.g. to produce subgraphs for ML training, run SPARQL queries, etc.). To ensure that the code remains functional and to detect breaking changes in data from upstream sources, we run our pipeline regularly using a continuous integration system^30^. This pipeline emits a KG that integrates all available data sources, in both TSV and RDF format, and also loads this KG into a Blazegraph database. A YAML file containing an inventory of the Biolink categories and Biolink associations of all data in the KG is also produced during the merge step (Figure 1).

### Generation of training and test edges for ML applications

To generate positive edges, a set of positive test edges equal in number to [(1 - train_fraction) * number of edges in input graph] is randomly selected from the edges in the input graph, where train_fraction is a number between 0 and 1 indicating the fraction of the graph to use for training. Positive test edges are selected such that removing them from the graph would not break it into disjoint components. These positive edges are removed from the edges of the input graph and are then emitted as the training edges. A set of negative edges is constructed by randomly selecting pairs of nodes that are not connected by an edge in the input graph. The number of negative edges emitted is equal to the number of positive edges emitted above. If the user requests a validation set, the positive test edges are divided equally to yield positive test and validation sets, and negative test edges are divided equally to yield negative test and validation sets.

### Embeddings and t-SNE plot for knowledge graph visualization

We generated embeddings from our KG using Embiggen^31^, our Python library for graph embedding and machine learning, using node2vec with a skip-gram model, 128 embedding dimensions, and parameters *p* and *q* of 1 (which are typically used default parameters for node2vec)^32^. These embeddings were used to generate a t-SNE plot that represents the embeddings for each node in two-dimensional space, using MulticoreTSNE^33^ (Figure 6).

## DISCUSSION

### A ‘KG-hub’ pattern for data sharing

The pattern used in the KG-COVID-19 framework as described in Figure 1 may be generally useful for data sharing among scientific communities. In the KG-COVID-19 framework, each data source is transformed and output as a separate graph, which is later combined with graphs for other data sources according to the needs of the user. Although the subgraphs from the various data sources (e.g., STRING, Drug Central) are produced locally by KG-COVID-19, our framework could easily consume and incorporate graphs generated by other members of the community. The exchange of data via a ‘KG-Hub’ would eliminate the duplication of effort that occurs when researchers separately transform and prepare data, and might also facilitate the formation of a data sharing portal for easier exchange of data.

### Comparison with similar projects

There have been a few parallel efforts to construct KGs to integrate COVID-19 data, each integrating different data sources and constructed for different purposes. Several efforts have constructed KGs by ingesting and transforming scientific literature,^34,35^ some with a few additional types of data also included, such as confirmed case and mortality data;^36^ clinical information, drug trial, and sequencing data;^37^ drug, drug trial and genome sequence data;^38^ diseases, chemicals, and genes^39^. Other KG efforts ingest a wider array of data, including diseases, genes, proteins and their structural data, drugs, and drug side effects;^40^ pathways, proteins, genes, drugs, diseases, anatomical terms, phenotypes, microbiome;^41^ genes, proteins, diseases, phenotypes, genome sequences;^42,43^ geographic, viral genes, genes and proteins.^44^ Several projects have focused specifically on integrating a wide variety of COVID-19 data to create KGs to investigate drug repurposing.^45–47^ The effort described here is unique in that it allows users to more flexibly remix specific data types from specific data sources (by virtue of its use of the KGX tool), it integrates more tightly with ontologies (HPO, Mondo, and GO) and with downstream machine learning tools (i.e. Embiggen), it offers a more detailed summary of the contents of its KG in a machine readable format, it covers a wider range of input data sources, and it automatically incorporates new and updated data.

### ID normalization challenges for SARS-CoV-2 entities

Since the usefulness of a KG depends on its connectedness, ID normalization is crucial. Normalization of IDs for SARS-CoV-2 entities in particular is challenging, for several reasons. First, SARS-CoV-2 produces identical cleavage products from different polyproteins, and UniProt assigns a different ID to each of these identical cleavage products. For example, UniProt uses PRO_0000338259 to identify the cleavage product nsp5, the 3C-like protease, if it is cleaved from replicase polyprotein 1a, and PRO_0000449623 if it is cleaved from replicase polyprotein 1ab. Protein Ontology, in contrast, uses PR_000050274, irrespective of the polyprotein from which it was cleaved. Note that the UniProtThe “PRO_” prefix is unrelated to the Protein Ontology namespace. For our KG, it is crucial that identical proteins be represented with a single node such that other information can be efficiently linked to them. We arbitrarily chose PRO_0000449623 as the ID to represent this cleavage product, and all other IDs for this cleavage product are stored as cross references for this node in our KG. Second, each cleavage product can have a large number of synonyms. For example, nsp5 has at least 40 synonyms that are used in the literature (e.g., 3CL-PRO, 3CLp, Mpro, 3C-like proteinase). Furthermore, some synonyms (e.g. ‘S’ for spike protein) are difficult to recognize when applying NLP to SARS-CoV-2 literature, which represents a further challenge for computationally identifying the occurrences of such entities in text. We have compiled our canonical IDs, synonyms, and cross references for each SARS-CoV-2 protein and cleavage product in our KG in a publicly available file in GPI format: https://github.com/Knowledge-Graph-Hub/kg-covid-19/blob/master/curated/ORFs/uniprot_sars-cov-2.gpi

### Conclusion

Knowledge graphs provide a way of integrating heterogeneous data from different sources and combining different data modalities. KG-COVID-19 generates a KG for COVID-19 focused around molecular and chemical information, and enables complex queries over relevant biological entities as well as machine learning to generate graph embeddings for making predictions. The lightweight framework we have developed provides a rapid route for bringing together new sources of data and knowledge, including KGs from several different sources, to form a “hub” to support COVID response efforts..

## DATA AND CODE AVAILABILITY

The Python code for KG-COVID-19 and the knowledge graph containing all data sources (in RDF and TSV format) are freely available at the KG-COVID-19 project wiki: https://github.com/Knowledge-Graph-Hub/kg-covid-19/wiki

The Python code is distributed under a BSD3 license.

A SPARQL endpoint is here:

http://kg-hub-rdf.berkeleybop.io/blazegraph/#query

## ACKNOWLEDGMENTS

This work was supported by grants from the Director, Office of Science, Office of Basic Energy Sciences of the U.S. Department of Energy [to J.R., D.U., S.C., N.L.H., M.J., C.J.M], the Laboratory Directed Research and Development (LDRD) Program of Lawrence Berkeley National Laboratory under U.S. Department of Energy Contract No. DE-AC02-05CH11231, the NIH (Monarch R24 OD011883, Illuminating the Druggable Genome U01 CA239108-01), a Training Grant from the NLM, NIH to the University of Colorado Anschutz Medical Campus Computational Bioscience Training Program [T15LM009451 to T.J.C.], the National Virtual Biotechnology Laboratory (NVBL), and the Google Cloud COVID-19 Research Grants program.

## AUTHOR CONTRIBUTIONS

The KG-COVID-19 framework was conceived and designed by J.R., D.U., M.P.J., C.J.M, T.J.C., N.M., S.C., V.R., and P.N.R., software was written by J.R., D.U., L.C., T.F., M.P.J., and the manuscript was prepared by J.R., D.U., M.P.J., C.J.M., H.B., N.H., M.M.T.

