## Supplemental Figures 1 and 2 for "KG-COVID-19: a framework to produce customized knowledge graphs for COVID-19 response"

**SUPPLEMENTAL INFORMATION DESCRIPTION**

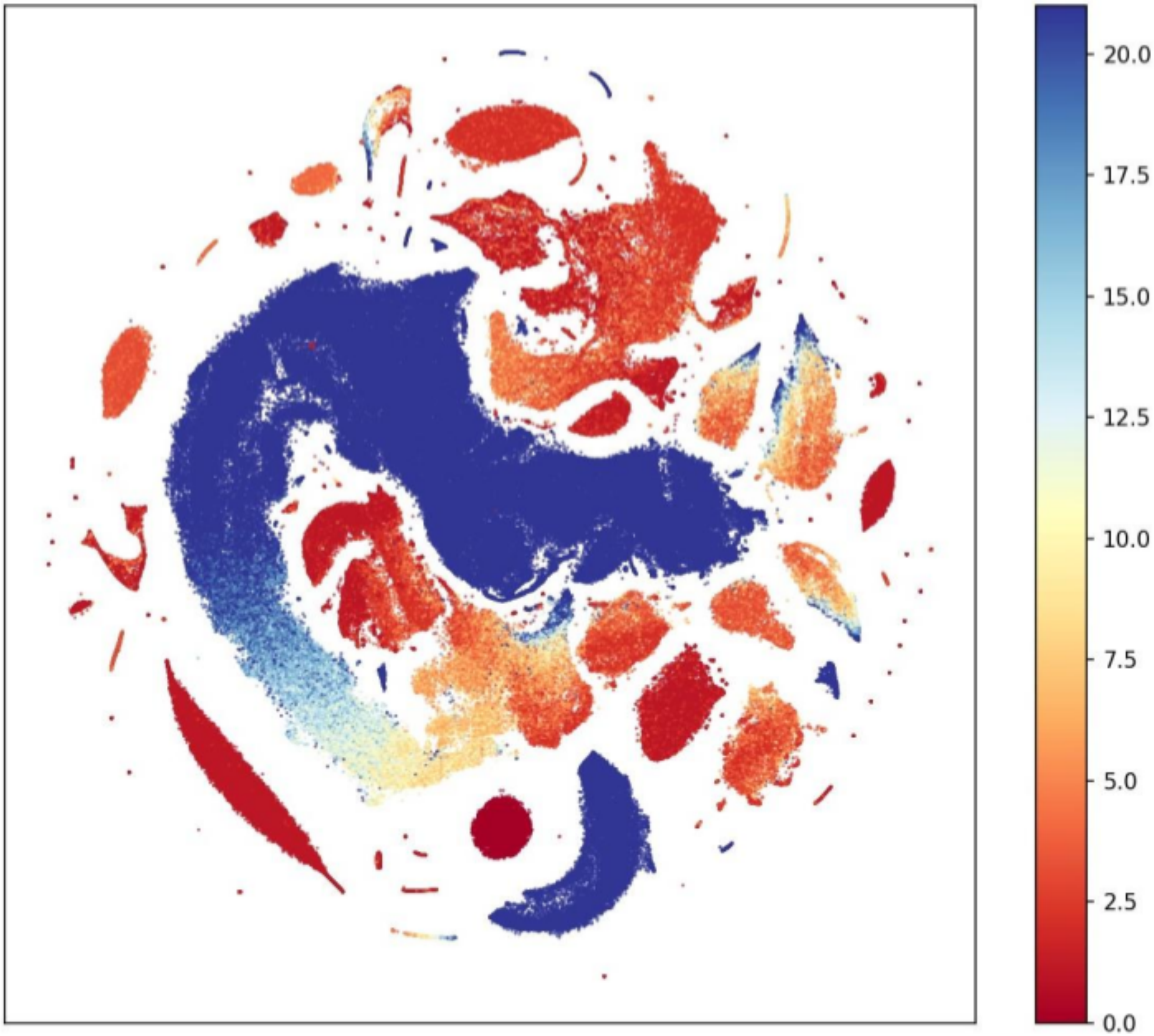


Supplemental Figure 1. Visualization of Biolink categories of nodes in the KG-COVID-19 knowledge graph by node degree. Embeddings were created for each node in the KG-COVID-19 knowledge graph and t-SNE was performed as described in Experimental Procedures. Colors indicate the degree for each node. Maximum node degree displayed is 21 (3 times the median node degree), and nodes with a higher degree were set to 21.

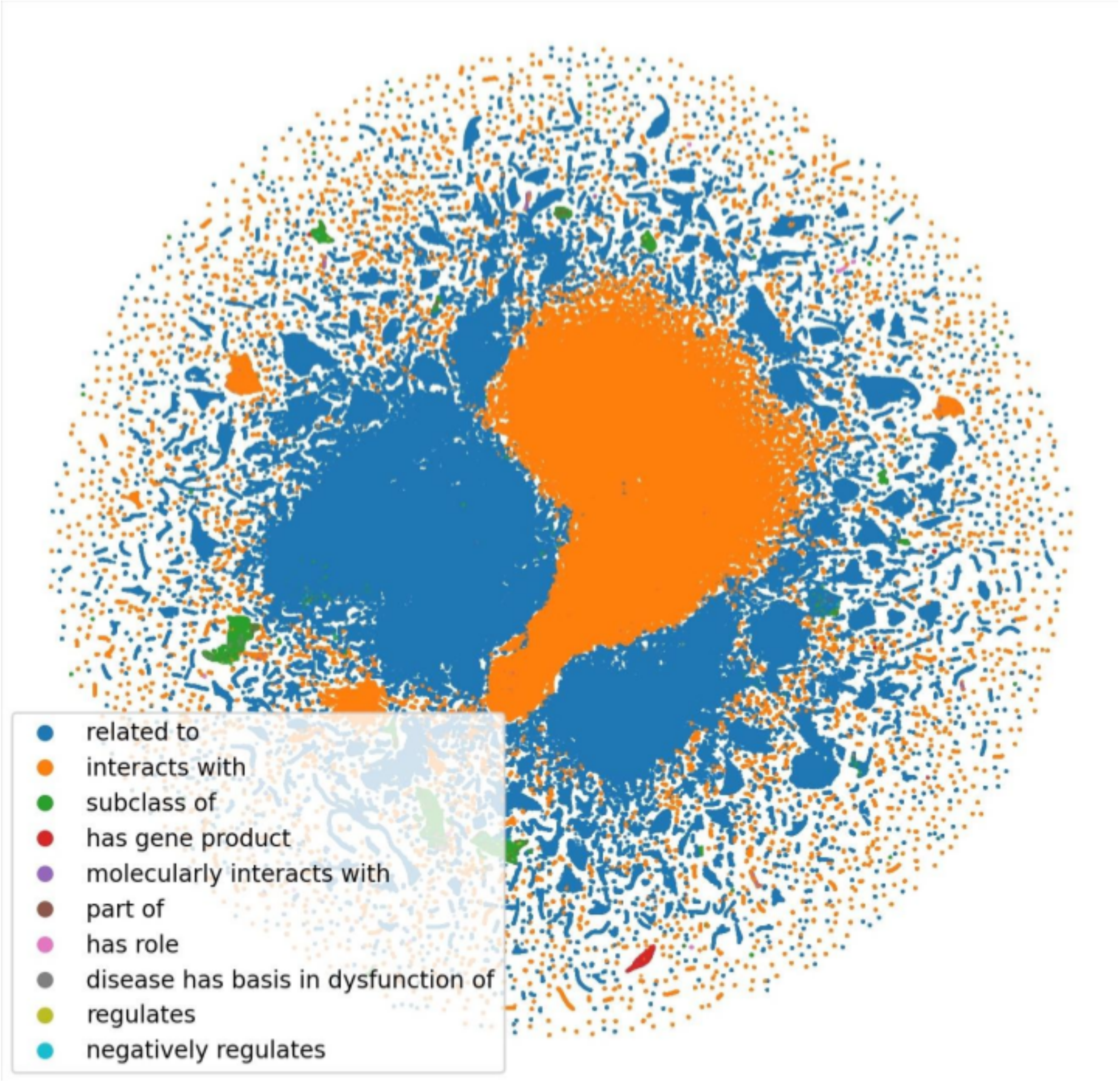


Supplemental Figure 2. Visualization of Biolink association type of edges in the KG-COVID-19 knowledge graph. Embeddings were created for 30% of edges in the KG-COVID-19 knowledge graph, sampled randomly, using Hadamard as edge embedding method and t-SNE was performed as described in Experimental Procedures. Edges categorized with one of the ten most numerous Biolink associations were then selected. Colors indicate the Biolink association type for each node.
